# MCT4-Associated Lactate Export Defines Myeloid Metabolic Reprogramming during Head and Neck Cancer Progression

**DOI:** 10.64898/2026.10.13.758108

**Authors:** Thomas Tilton, Young Min Park, Haiqi Zhu, James Paolillo, Chi Zhang, De-Chen Lin

## Abstract

Head and neck squamous cell carcinoma (HNSCC) development is accompanied by metabolic remodeling of the tumor microenvironment, yet how metabolic programs evolve across specific cell populations during tumor progression remains poorly understood. We integrated single-cell RNA-sequencing data from five HNSCC cohorts comprising 645,351 cells from 114 patients and inferred metabolic flux using single-cell Flux Estimation Analysis (scFEA).

Myeloid cells showed the highest and most consistent metabolic allocation to lactate export among major cell types. In ten patients with paired normal mucosa and carcinoma, myeloid lactate-export allocation increased from 1.72% to 3.31% of total inferred metabolic flux, with an increase in every patient. SPP1+ macrophages showed the highest lactate-export allocation in carcinoma.

Lactate export was predominantly associated with MCT4/*SLC16A3*. In carcinoma, myeloid *SLC16A3* expression exceeded *SLC16A1* by 17-fold, and CPTAC-HNSCC confirmed tumor enrichment of MCT4 protein. Together, these findings identify increased MCT4-associated lactate export as a prominent feature of myeloid metabolic reprogramming during HNSCC progression.

## Introduction

Tumors accumulate lactate as an outcome of the Warburg effect, where they utilize glycolysis to produce ATP and intermediate metabolites, even in aerobic conditions [1]. Aerobic glycolysis accounts for over half of a cancer cell’s ATP production [2]. Compared with normal blood lactate concentrations (0.5–2.2 mM), lactate concentration in the tumor microenvironment (TME) has been reported to reach 10–40 mM [1]. The increased lactate production results in transcription and metabolism changes that enhance malignant transformation and acidifies the TME, which further modulates cellular interactions [1].

Lactate is produced during the conversion of pyruvate to lactic acid by lactate dehydrogenase (LDHA). Lactate is the deprotonated version of lactic acid that exists at a neutral pH. As lactate accumulates, cells rely on monocarboxylate transporters (MCT1 and MCT4 in particular) to secrete lactate from the cytoplasm to extracellular fluid, resulting in an extracellular pH drop. Lactate also activates tumor-promoting genes, including vascular endothelial growth factor (VEGF) for neoangiogenesis, transforming growth factor beta (TGF-beta), interleukin-1 (IL-1), and HIF-1, even in normoxic tumor cells. Because HIF-1 regulates the shift toward glycolysis, lactate can feed back to enhance the process that produces it [2].

The MCT4 protein, encoded by the *SLC16A3* gene, has been implicated in oral carcinoma. Bisetto et al. reported that MCT4 was overexpressed in oral premalignant lesions and carcinoma by immunohistochemistry in nine human tumors and in a 4NQO mouse model of oral carcinogenesis [3]. However, these analyses could not resolve whether this increase reflects a shift in myeloid cell composition or metabolic reprogramming within myeloid populations.

We used single-cell data from HNSCC patients with paired normal and tumor tissues to address this gap. Using a metabolic flux inference algorithm, scFEA [4], we estimated flux for 168 metabolic modules from single-cell transcriptomes. Because scFEA-derived flux magnitudes are not directly comparable across cell types, we focused on metabolic allocation, defined as the fraction of a cell subtype’s total inferred metabolic flux assigned to each metabolic module. This approach captures changes in the relative distribution of metabolic activity across pathways, while remaining insensitive to uniform changes in overall metabolic activity. Allocation also allows comparison across different cell types. Given that scFEA estimates represent model-inferred rather than directly measured metabolic flux, we further applied stringent criteria to identify robust and interpretable changes. This analysis allowed us to determine how individual cell populations redistribute their metabolic activity during progression from normal oral mucosa to HNSCC.

## Methods

### Data and harmonization for 5-cohort single-cell atlas

Five published head and neck single-cell datasets were assembled, comprising 645,351 cells from 114 patients provided as 167 samples: He [5], Kim [6], Koh [7], Pittet [8], and Tirosh [9]. Datasets are referred to by their senior author. Per-cohort composition by tissue class is shown in Figure 1.

**Figure 1.**
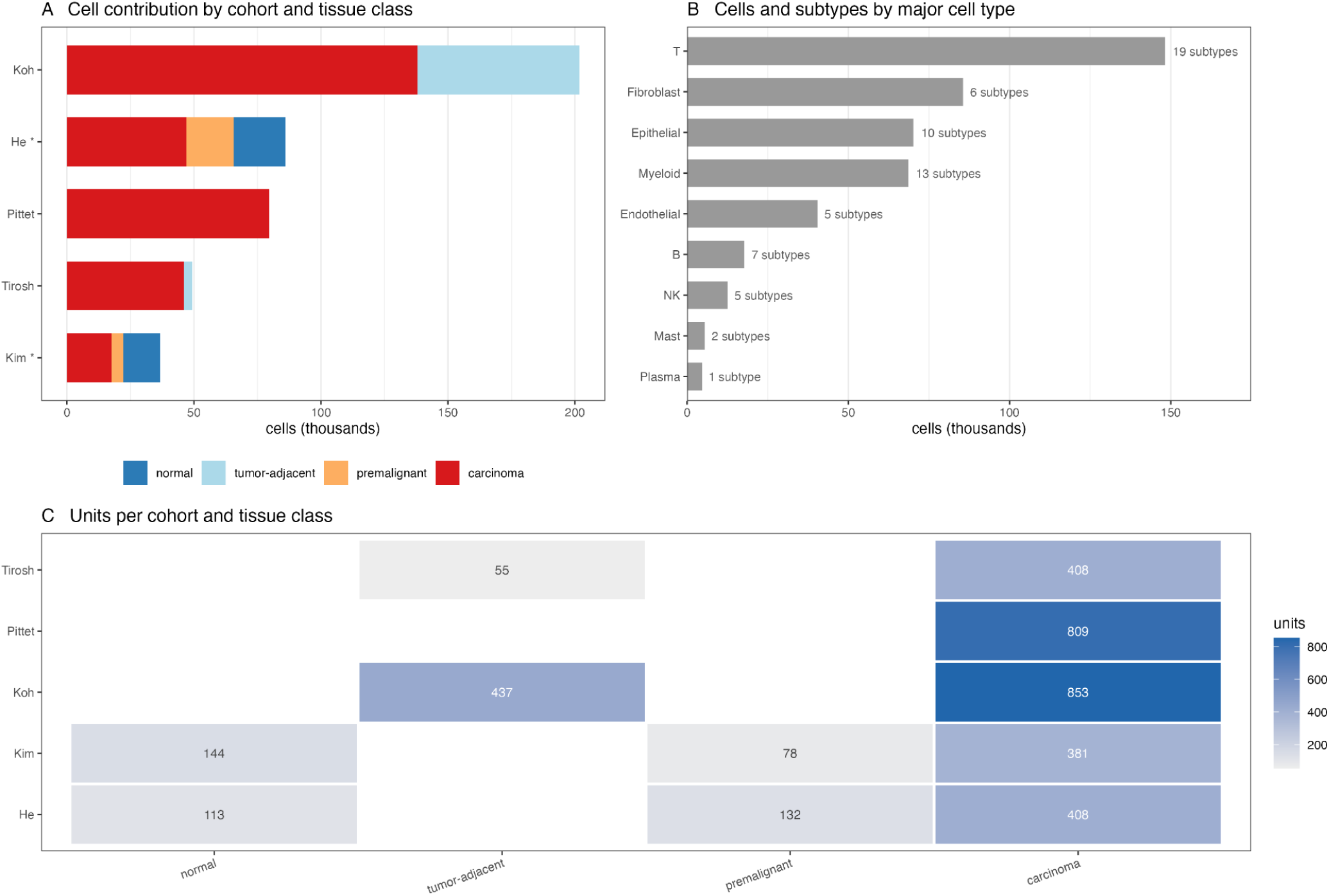
HNSCC single-cell atlas and analytical framework. (**A**) Cell counts by cohort and tissue type; asterisks indicate cohorts with paired normal mucosa and carcinoma used for progression analyses. (**B**) Cell counts and number of subtypes by major cell type. (**C**) Analytical units, defined as patient × tissue × cell subtype combinations containing ≥10 cells, by cohort and tissue class. Blank entries indicate combinations that were unavailable or insufficiently represented.

Cell type annotation was harmonized to nine major cell types (annot_4th) and 62 cell subtypes (finaltype). Tissue was classified as distant normal mucosa (NL), tumor-adjacent (NL-contact), premalignant (Pre, LP) or carcinoma (CA and subtypes). Distant normal and tumor-adjacent are reported separately throughout. The premalignant class pools two labels used by different cohorts, Pre from He and leukoplakia (LP) from Kim, but progression analysis was run within each cohort, so a given test uses only one of these labels. Lymph node, recurrent, and metastatic tissue were excluded from all analyses.

Cohorts were included in specific analyses based on sample availability. Because only the He and Kim cohorts contained normal mucosa, premalignant lesion and carcinoma, and paired normal mucosa and carcinoma from the same patients, analyses of metabolic changes during tumor progression were restricted to these two cohorts. All five cohorts were included in analyses conducted within carcinoma, including the comparison of metabolic allocation between major cell types, the ranking of myeloid subtypes, the empirical null and the evaluation of flux-enzyme concordance. Although the Koh and Tirosh cohorts also contained tumor-adjacent tissues, these samples were not included in the present analyses because they are biologically distinct from normal mucosa and therefore were not considered appropriate substitutes for the paired normal tissues used in the progression analysis.

### Unit of analysis and filtering criteria

Analyses were performed on pseudobulk units, each defined as a unique combination of patient, tissue, and cell subtype. Aggregating in this manner ensures a patient contributes to a single observation per tissue and subtype rather than one per cell, thereby preventing samples with greater numbers of sequenced cells from disproportionately influencing the analysis.

Units were required to contain at least 10 cells. The robustness of this threshold was evaluated across alternative cutoffs. A threshold of 10 yielded identical analytical coverage to a threshold of 8, recovering the same thirteen progression comparisons, while removing 350 more degenerate units. A threshold of fifteen reduced analytical coverage, eliminating two progression comparisons. The same threshold was applied to every analysis and to both pseudobulking strategies (see Methods, Flux estimation), to ensure comparability.

After filtering, the atlas contained 5,459 units. Of these, 3,818 were from the oral axis (normal mucosa, tumor-adjacent tissue, premalignant lesion and carcinoma) and the remainder from lymph node, recurrent and metastatic samples, which are excluded from analyses reported here.

Additional sample-size criteria apply to specific analyses. Comparisons required at least three patients per group, and at least five patients per arm for HPV-stratified comparisons. For correlations calculated independently within each source cohort and subsequently combined, a cohort had to contain at least eight units for the given cell type and metabolic module, thereby limiting unstable correlation estimates arising from small sample sizes.

NK cells were insufficiently represented in the primary analysis, with a median of two cells per unit before filtering and approximately 75% of units falling below the 10-cell threshold. Consequently, results for NK cells in the primary analysis should be interpreted cautiously, as the absence of significant findings may reflect limited sampling. This limitation does not apply to the secondary analysis, in which flux was estimated at the single-cell level before aggregation.

### Imputation

Gene expression was imputed with MAGIC before flux estimation, in accordance with the imputation procedure in the scFEA pipeline. MAGIC was run independently to the normalized expression matrix of each major cell type, and the same imputed matrices were used for both pseudobulking strategies. Thus, the two analytical strategies differ only in the order of aggregation and flux estimation.

All MAGIC parameters were at default values (knn = 5, decay = 1, t = ‘auto’, n_pca = 100), consistent with the internal imputation procedure implemented by scFEA. The exact solver was used for all nine major cell types. The MAGIC neighborhood graph was built on expression from all genes across the full normalized matrix. At the imputation step within MAGIC, only the 661 genes belonging to scFEA’s metabolic modules were imputed and returned, which reduces computation without altering the graph. scFEA was run with --sc_imputation False in both arms to avoid repeated imputation.

Imputation was performed at the single-cell level, before units are formed. Pseudobulk values are therefore averages of imputed single-cell values rather than imputed averages. Because downstream quantities derived from these values are influenced by MAGIC smoothing, differential expression analyses were instead performed using raw counts.

### Flux estimation

Metabolic flux was estimated using scFEA against 168 metabolic modules (M1-M168) of the M171 metabolic network encoded in the method. The algorithm was implemented using its default settings, but minor implementation modifications were made to enable analysis of large datasets and GPU-based computation. The implementation was modified to add --mode {train, predict} and --model_file. This allowed us to train a model on a smaller subset of samples, and then predict with the trained model on the entire atlas. For GPU compatibility, we called .cpu() before .numpy() because PyTorch tensors on a GPU cannot be converted directly to NumPy arrays, so .cpu() moves the tensors to CPU memory. Hyperparameters for scFEA were the code defaults: LAMB_BA 1, LAMB_NG 1, LAMB_CELL 1, LAMB_MOD 1e-2, LEARN_RATE 0.008, torch.manual_seed(16), and --train_epoch 100, set explicitly in every submission script and identical across both pseudobulking arms. The scFEA source used was commit 4c1fb76d52f07bafad84ce7686ad7c3acfcf0126 (9 November 2022).

scFEA was run independently for each major cell type (on pseudobulk units in the primary analysis and on single cells in the secondary analysis; see below), with all cohorts and all tissue types for a given cell type included within the same model. Thus, model parameters were shared across conditions and cohorts within a cell type, but not across cell types. Because scFEA normalizes flux estimates relative to mean expression within each run, absolute flux magnitudes are not directly comparable across cell types. We therefore used metabolic allocation, the proportion of total inferred flux assigned to each module, for cross-cell-type comparisons.

To assess the robustness of flux estimates to the order of aggregation and inference, two complementary pseudobulking strategies were implemented. In the primary run, gene expression was aggregated via pseudobulk first, followed by scFEA flux estimation. In the secondary run, scFEA was first run at the single-cell level, followed by pseudobulk aggregation of the resulting flux estimates. The primary and secondary strategies differ in whether metabolic network constraints are applied to individual cells or aggregated expression profiles. We therefore considered findings robust only when they were supported by both analytical strategies.

Flux values close to zero occasionally yielded small negative values, a residue of numerical optimization rather than a meaningful quantity; modules where this affected a summary statistic were excluded from interpretation. This occurred more frequently in the secondary analysis (40 of 168 modules) than in the primary analysis (6 of 168 modules). Accordingly, the primary analysis was used for quantitative reporting, whereas the secondary analysis served as a robustness analysis.

### Metabolic allocation

Metabolic allocation was defined as a specific metabolic module’s flux divided by the unit’s total flux (unit = patient × tissue × cell subtype) across all usable modules, expressed as a percentage. Thus, allocation reflects the relative distribution of metabolic activity across pathways, with module allocations summing to 100% within each unit. A uniform change in flux across all modules would therefore not alter metabolic allocation.

### Empirical null and flux-enzyme concordance

scFEA infers metabolic flux from gene expression under metabolic network constraints. As a result, inferred flux may not necessarily track expression of the genes assigned to each module. We therefore used flux–enzyme concordance as a quality-control criterion to identify module–cell type combinations suitable for interpretation. Because gene expression is an input to scFEA, this criterion was used only for filtering and not as independent validation of inferred flux.

For each module, concordance was defined by the gene showing the strongest correlation with inferred flux. We used the strongest rather than the average correlation because only a subset of genes within a module may contribute substantially to the corresponding reaction in a given cell type. For example, M_6 contains LDHA, LDHAL6A, and LDHAL6B. M_6 flux correlated strongly with LDHA (ρ = 0.79), whereas its correlation with mean module expression was substantially lower (ρ = 0.18). This analysis ensures that the reported flux changes can be supported by the expression change of at least one enzyme involved in the reaction. Concordance was evaluated per major cell type and module, matching the resolution at which scFEA models were trained. Comparisons from the same major cell type and module therefore share a concordance value.

Because selecting the strongest-correlating gene favors modules containing more genes, we generated a matched empirical null that accounts for module size. For a module containing *k* evaluable genes, *k* correlations were sampled from correlations between module flux and non-module metabolic genes within the same cell type, and the maximum was retained. This procedure was repeated 20,000 times, and the 99.9th percentile of the resulting distribution was used as the matched empirical null threshold for that module and cell type.

Single-gene modules were excluded because concordance between inferred flux and its sole input gene cannot provide an informative quality-control measure. Modules lacking sufficient observations for correlation analysis were also excluded. Among 162 usable modules, 28 were single-gene modules and six lacked sufficient observations, leaving 128 modules and 304 evaluable cell type–module combinations. Of these, 44 exceeded their matched empirical null thresholds and were retained for biological interpretation.

### Statistical analysis

Progression was tested in two ways: a module-wide trend analysis across all cell subtypes and modules, and paired comparisons within patient for the results reported in detail. For the module-wide analysis, allocation was tested for a monotonic trend across the three ordered tissue classes: normal mucosa, premalignant lesion, and carcinoma. We used the Jonckheere–Terpstra test with the normal approximation, within each cohort separately. A comparison was included only if that cell subtype had units in all three classes, with at least two per class. Cohort-specific z-scores were combined by Stouffer’s method, weighted by the square root of the number of patients in each cohort, and the combined p-values were adjusted by the Benjamini–Hochberg procedure across all 2,268 cell subtype–module comparisons in that pseudobulking run. Statistical significance was defined by FDR < 0.10 alone.

Two additional criteria were applied to prioritize statistically significant comparisons for biological interpretation; these are filters, not tests of significance. First, an effect-size threshold was set at the 99th percentile of the observed effect-size distribution for that run, so that only the largest 1% of changes were carried forward. The effect size for each comparison was the mean of the two cohorts’ effect sizes, weighted by patient count, and the percentile was computed over all 2,268 comparisons before FDR filtering. The 99th percentile was a stringency choice rather than a calibrated threshold, and was derived empirically rather than fixed in advance to accommodate differences in effect-size distributions arising from cell-type-specific scFEA modeling and the two pseudobulking strategies. Second, effects were required to have the same direction in both cohorts, so that combined significance was supported by consistent effects across the two independent cohorts.

The second approach was used for the comparisons reported in detail: myeloid export allocation between normal mucosa and carcinoma, the nested subtype sets, and the between-cell-type comparisons. These comparisons were paired within patient and tested using the Wilcoxon signed-rank test. With ten nonzero paired observations, the minimum attainable two-sided exact P value is 0.00195, corresponding to all ten paired differences occurring in the same direction; 93 of 162 myeloid modules reached this minimum. Because the small sample size limits the resolution of exact P values, effect sizes were evaluated separately. Medians are reported for allocation because the distributions are bounded and right-skewed.

HPV-stratified comparisons were tested within carcinoma, within each cohort, by the Wilcoxon rank-sum test on patient-level values, requiring at least five patients per HPV arm. Cohort z-scores were combined by Stouffer’s method, weighted by the square root of the smaller arm, across 4,857 cell subtype–module comparisons. A significant comparison was retained only if the module’s flux shifted by at least 10% and the effect was in the same direction in both cohorts; comparisons testable in only one cohort were reported but could not meet the concordance criterion.

Multiple testing was controlled by the Benjamini–Hochberg procedure within each analysis, at a false discovery rate of 0.10.

All statistical analyses were performed in R version 4.4.1. Wilcoxon signed-rank and rank-sum tests used wilcox.test and multiple testing correction used p.adjust, both from the stats package. The Jonckheere-Terpstra trend test with the normal approximation and the Stouffer combination of z-scores were implemented directly rather than through a package. Cox proportional hazards models and the cox.zph diagnostic used the survival package version 3.8-3.

### Differential expression analysis

Differential expression was computed on patient-level pseudobulk expression profiles defined by tissue and major cell types. Raw gene counts were summed across cells within each patient–tissue–cell type profile, giving one count profile per patient per disease stage per cell type. Profiles were required to contain at least ten cells. Genes were filtered with edgeR’s filterByExpr (version 4.2.2), counts were normalized by the trimmed mean of M-values, and differential expression was tested with a quasi-likelihood F test under a paired design (∼ patient + stage), so that each patient serves as their own control.

These profiles use raw counts from the source data rather than the MAGIC-imputed metabolic gene panel supplied to scFEA, so the differential expression results do not depend on the imputation step.

The differential expression analysis was restricted to the patients used in the flux analysis, to maintain consistent patient sets across metabolic and transcriptional comparisons. This resulted in 10 paired patients for myeloid cells, 8 for fibroblasts, and 9 for T cells, reflecting differences in the number of patients meeting cell-number requirements for each cell type. After filtering, 12,028 genes were tested in myeloid cells, 16,947 in fibroblasts and 12,153 in T cells.

### Survival analysis

We performed survival analysis using the TCGA-HNSC cohort accessed through cBioPortal [13,14]. Expression values for each gene were log2-transformed and z-scored across samples. The lactate metabolic program score is the mean z-score of SLC16A3, LDHA and HK2. Overall survival was modeled by multivariable Cox proportional hazards, adjusted for age at diagnosis and stratified by stage (I-II versus III-IV), with hazard ratios reported per standard deviation. Patients missing age or stage were excluded, leaving 417 patients and 184 events of the original 514 with survival data. The proportional hazards assumption was assessed with scaled Schoenfeld residuals and was not violated (global P = 0.49). We used a median split for the Kaplan-Meier curve in Figure 6A for visualization purposes only. All hazard ratios and P values we report derive from the continuous lactate metabolic program z-score, not the median split.

### External datasets

Two independent HNSCC single-cell RNA-sequencing cohorts, not part of the five cohort atlas, were used for the lactate transporter comparison: Cillo (GSE139324, restricted to tumor-infiltrating cells) [10] and Kürten (GSE164690, tumor only) [11]. For consistency, analyses of our integrated atlas were similarly restricted to carcinoma samples.

Proteomic data are from CPTAC-HNSCC [12], accessed through the cptac Python package (v1.5.14), using proteomics from the umich pipeline (182 samples × 12,224 proteins). Protein abundance was compared between 116 tumor and 66 normal-adjacent samples with the Mann-Whitney U test; the reported difference is the tumor median minus the normal-adjacent median in that pipeline’s normalized units. This comparison is unpaired.

Bulk expression and clinical data from TCGA-HNSC, including survival and precomputed hypoxia scores, were accessed through cBioPortal [13,14].

### Ethics statement

This study did not involve new human subjects, specimens, or animal experimentation. Only previously published, de-identified data were used, and the research was conducted in accordance with the principles of the Declaration of Helsinki. Ethics approval and informed consent were obtained by the investigators of each source study, from an institutional review board or equivalent ethics committee, and are reported in the original publications [5–9]. Data under accession HRA004648 were obtained through an approved data access agreement. The remaining datasets, GSE181919, GSE198315, GSE234933, GSE182227, GSE139324, GSE164690, TCGA-HNSC, and CPTAC-HNSCC, are publicly available in de-identified form. Because this work analyzed only de-identified data that are not individually identifiable, it did not constitute human subjects research and no additional institutional review board approval was required.

### Code Availability

Analysis code will be available after publication.

### Data Availability

The five source single-cell datasets are publicly available under accessions HRA004648 (He), GSE181919 (Kim), GSE198315 (Koh), GSE234933 (Pittet), and GSE182227 (Tirosh). HRA004648 is controlled access and readers should apply to that repository directly. The two external validation cohorts are GSE139324 (Cillo) and GSE164690 (Kürten). Proteomic data are from CPTAC-HNSCC, accessed through the cptac Python package. Survival and expression data are from TCGA-HNSC via cBioPortal. Data objects will be made available upon official publication.

## Results

### HNSCC single-cell atlas and analytical framework

We assembled 645,351 cells from 114 patients across five published cohorts (He, Kim, Koh, Pittet, Tirosh; Fig. 1A) with harmonized annotation resolving to nine major cell types and 62 cell subtypes on the oral axis (68 subtypes across all tissue; Fig. 1B). The unit of analysis was a pseudobulk profile representing a specific cell subtype and tissue type from an individual patient (patient × tissue × cell subtype). We set a threshold requiring a minimum of ten cells for each possible combination to pass filtering. Of 8,639 units created (total number of distinct patient × tissue × cell subtype across all tissue), 5,459 met the 10 cell threshold.

Our analyses focused on the oral disease continuum: normal mucosa, tumor-adjacent tissue, premalignant lesion, and carcinoma. These disease states were characterized by 3,818 units from 82 patients. Lymph node, recurrent, and metastatic tissue were excluded from our analyses. Metabolic flux was inferred independently for each major cell type using scFEA, with all cohorts and tissue states included within each cell-type analysis.

To examine metabolic changes associated with progression from normal mucosa to carcinoma, we focused on the He and Kim cohorts, the two datasets containing matched normal and carcinoma tissues from the same patients. Ten patients had paired myeloid profiles meeting the inclusion criteria. Analyses performed within carcinoma incorporated data from all five cohorts.

### Metabolic allocation reveals selective pathway reprogramming during tumor progression

We first performed a systematic screen for metabolic changes across progression from normal mucosa through premalignant lesions to carcinoma. Statistical significance was assessed using a trend test across the three tissue classes, whereas effect size was defined by the change in allocation from normal mucosa to carcinoma. When absolute inferred flux was compared between normal mucosa and carcinoma across tested cell type–module comparisons, 66% showed increased flux in carcinoma, suggesting a broad increase in inferred metabolic activity (Fig. 2A). In contrast, analysis of metabolic allocation, which measures the proportion of total metabolic activity assigned to each metabolic pathway, identified increases in 38% and decreases in 62% of comparisons. Nineteen comparisons met the predefined FDR, effect-size, and cross-cohort concordance criteria (Fig. 2B), of which six additionally met the flux–enzyme concordance criterion and were retained for further downstream analyses (Fig. 2C). Additional candidate associations did not meet these criteria and were not pursued further (Supplementary Table S1). All nineteen survivors and their criterion status are shown in Supplementary Table S2.

**Figure 2.**
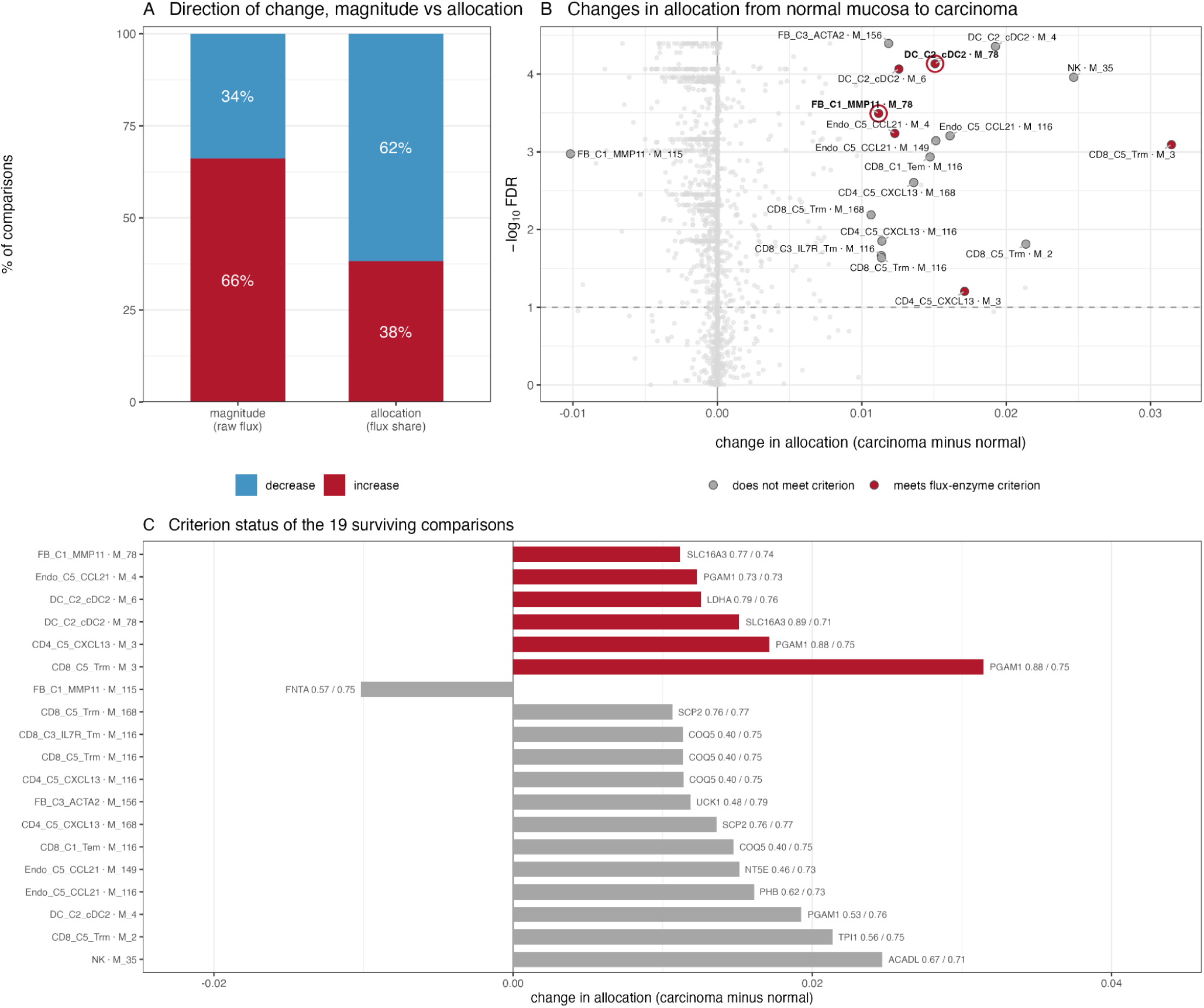
Metabolic allocation identifies pathway-specific changes during tumor progression. (**A**) Direction of metabolic changes based on absolute inferred flux versus metabolic allocation. (**B**) Meta-analysis of allocation changes across the He and Kim cohorts. Effect size is the change in allocation from normal mucosa to carcinoma; significance is from a trend test across normal mucosa, premalignant lesion, and carcinoma. Nineteen comparisons met the FDR, effect-size, and cross-cohort concordance criteria; six additionally met the flux–enzyme concordance criterion. Lactate-export modules are highlighted. (**C**) Flux–enzyme concordance for the 19 comparisons in (B). Labels give the best-correlating gene, its Spearman correlation with module flux, and the matched empirical null threshold (correlation / threshold). Red bars indicate comparisons whose correlation exceeded the threshold and were retained; grey bars did not.

### Myeloid cells preferentially allocate metabolism to lactate export

In carcinoma, myeloid cells showed the highest allocation to lactate export (M_78), with a median of 2.0% compared with 1.2% in T cells (Fig. 3A). In paired within-patient comparisons, myeloid allocation exceeded that of every other major cell type in 72–100% of patients. Among 67 patients with both populations represented, myeloid allocation exceeded T-cell allocation (the 2^nd^ highest) by a median of 1.09 percentage points (P = 1.5 × 10⁻¹¹).

**Figure 3.**
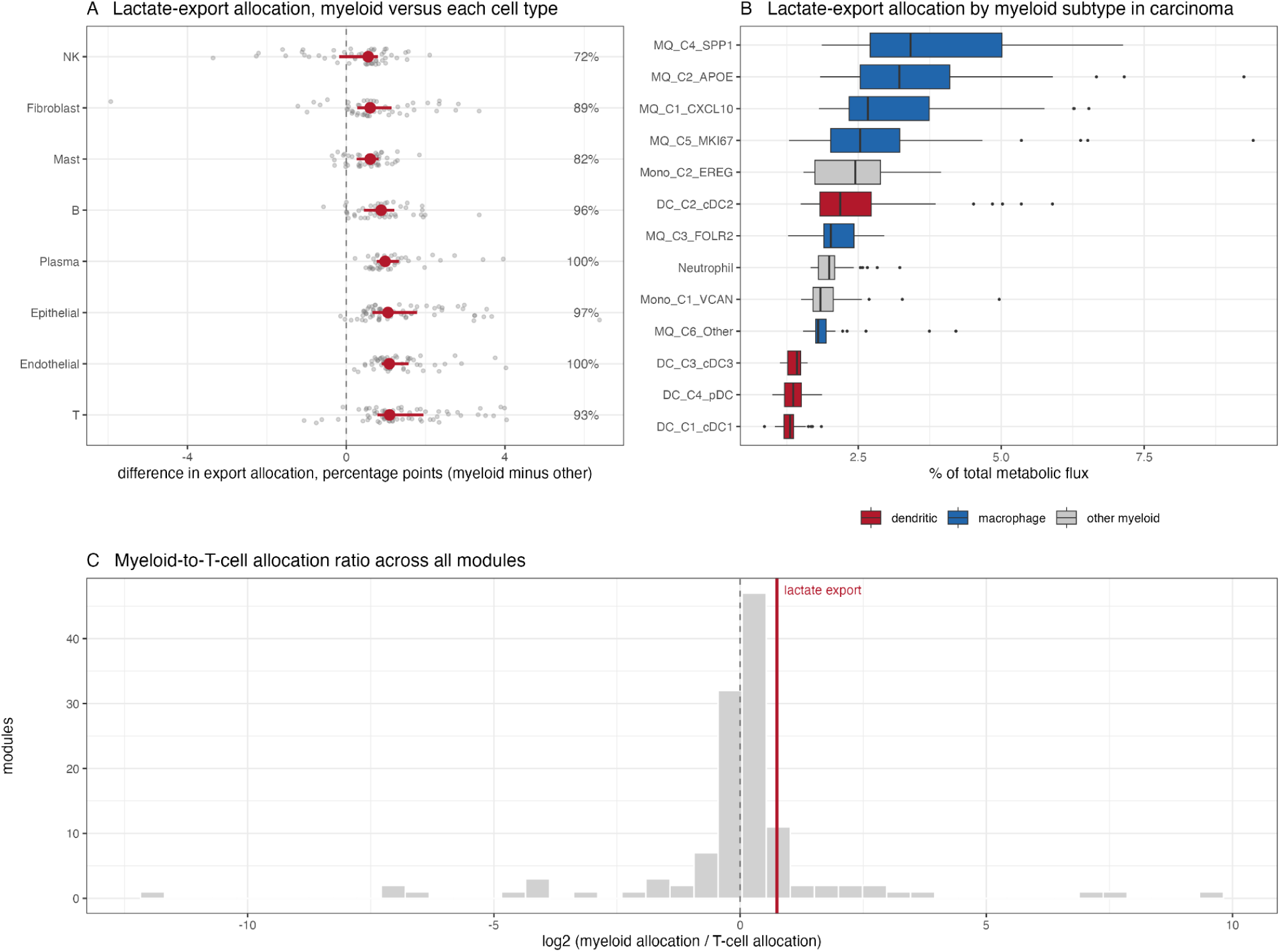
Lactate-export allocation across cell types and myeloid subtypes. (**A**) Lactate-export allocation across major cell types in carcinoma, compared within patients. Each point is one patient; the red dot and bar are the median and interquartile range. Percentages on the right are the share of patients with higher lactate-export allocation in myeloid cells. (**B**) Lactate-export allocation across myeloid subtypes in carcinoma (668 units). (**C**) Myeloid-to-T-cell allocation ratio across 125 evaluable metabolic modules; lactate export ranked at the 87th percentile.

Within the myeloid compartment, lactate-export allocation varied 2.6-fold across subtypes (Fig. 3B). SPP1+ macrophages showed the highest allocation (median, 3.4%), followed by APOE+ and CXCL10+ macrophages, whereas cDC1 and pDC populations showed the lowest (1.3%).

Notably, lactate export was distinguished more by the consistency than the magnitude of its myeloid enrichment. It ranked at the 87th percentile among 125 metabolic modules based on myeloid-to-T-cell allocation (Fig. 3C), but was preferentially allocated in myeloid cells across nearly all patients.

### Lactate-export allocation selectively increases in myeloid cells during tumor progression

In ten patients with paired normal mucosa and carcinoma, myeloid lactate-export allocation increased from a median of 1.72% to 3.31% of total metabolic flux, with an increase observed in every patient (P = 0.002; Fig. 4A). Among modules that increased in all ten patients, lactate export ranked second by effect size in both pseudobulking analyses and was the highest-ranked module meeting the flux–enzyme concordance criterion. The metabolic module ranked above it in each analysis (M_4 in the primary, M_116 in the secondary) did not meet the concordance criterion in either.

**Figure 4.**
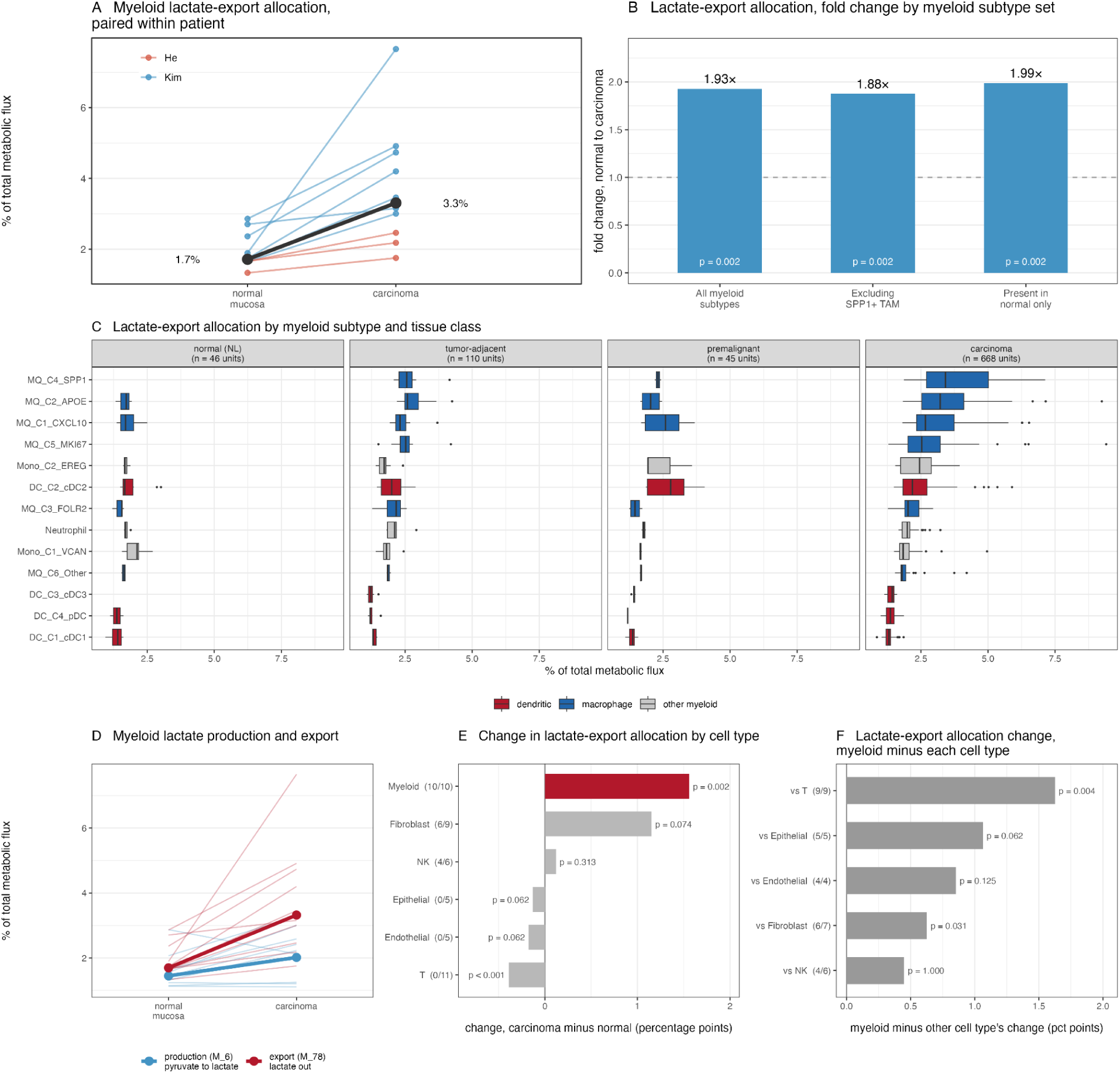
Lactate-export allocation preferentially increases in myeloid cells during progression from normal mucosa to carcinoma. (**A**) Paired myeloid lactate-export allocation in normal mucosa and carcinoma (n = 10). Lines represent individual patients; thick line indicates the median. (**B**) Paired analysis after excluding SPP1+ macrophages or restricting to myeloid subtypes represented in normal mucosa. (**C**) Lactate-export allocation across myeloid subtypes and tissue classes, ordered by carcinoma median. Empty cells indicate subtypes without evaluable units. (**D**) Paired changes in lactate export (M_78) and lactate production (M_6). (**E**) Changes in lactate-export allocation across major cell types with paired normal and carcinoma samples. Parentheses indicate patients with increased allocation/total patients evaluated. (**F**) Within-patient difference between changes in myeloid cells and each indicated cell type; positive values indicate a greater increase in myeloid cells.

This increase was not simply attributable to increased glycolytic activity. Allocation to lactate production (M_6; pyruvate to lactate) increased more modestly, from 1.46% to 2.18%, and in only 7 of 10 patients (P = 0.049; Fig. 4D). Moreover, M_8 (citrate to 2-oxoglutarate), a TCA-cycle module, also increased in all ten patients in both pseudobulking analyses, and met the flux–enzyme criterion in the primary analysis. Thus, increased lactate-export allocation was not accompanied by a corresponding reduction in TCA-cycle activity.

Importantly, increased lactate-export allocation from normal mucosa to carcinoma was preferential to the myeloid compartment (Fig. 4E). Myeloid allocation increased in all 10 paired patients (median change, +1.56 percentage points; P = 0.002), whereas T-cell lactate-export allocation decreased in all 11 paired patients (P = 0.001), with no significant increase in endothelial, epithelial, NK, or fibroblast populations. Patient counts differ between cell types because each requires sufficient cells of that type in both tissues. Comparing changes within the same patient, the myeloid increase exceeded that of every other cell type tested, including fibroblasts (6 of 7 patients, P = 0.031) and T cells (9 of 9, P = 0.004; Fig. 4F). Together, these findings identify increased lactate-export allocation as a selective feature of myeloid metabolic reprogramming during progression to carcinoma.

### Lactate-export reprogramming extends across myeloid subtypes

Three myeloid subtypes (SPP1+ macrophages, proliferating MKI67+ macrophages, and cDC3) had no evaluable units in normal mucosa (Fig. 4C). Because SPP1+ macrophages showed the highest lactate-export allocation in carcinoma, we asked whether their enrichment could account for the compartment-level increase. Across all myeloid subtypes, lactate-export allocation increased 1.93-fold from normal mucosa to carcinoma. The increase was largely unchanged after excluding SPP1+ macrophages (1.88-fold) or restricting the analysis to the 10 myeloid subtypes represented in normal mucosa (1.99-fold; Fig. 4B). All three comparisons were significant (P = 0.002) and reproduced in both pseudobulking analyses. Thus, increased lactate-export allocation is not driven solely by the emergence or enrichment of high-export myeloid subtypes in carcinoma.

We further examined whether a single myeloid subtype disproportionately contributed to this increase. Among subtypes with paired normal and carcinoma samples, none showed a significant change in lactate-export allocation relative to the overall myeloid compartment. However, the limited number of paired patients for individual subtypes (n = 4–8) precluded definitive subtype-level comparisons. Together, these findings support a broadly distributed increase in lactate-export allocation across the myeloid compartment rather than an effect dominated by a single subtype.

No lactate-handling module differed by HPV status in any myeloid subtype (minimum FDR = 0.85 across 24 comparisons in carcinoma; Supplementary Table S4). Most comparisons were testable in one cohort only, so this analysis is underpowered rather than negative, and HPV status is confounded with anatomic site in these data.

### MCT4 is the predominant lactate transporter expressed by myeloid cells

The lactate-transport module (M_78) comprises *SLC16A1* (MCT1), *SLC16A3* (MCT4), and *SLC16A4* (MCT5). In myeloid cells, M_78 flux correlated strongly with *SLC16A3* expression (ρ = 0.89), exceeding the matched empirical null threshold of 0.715, whereas *SLC16A1* showed substantially weaker correlation and *SLC16A4* was negatively correlated with inferred flux (Fig. 5A). *SLC16A3* accounted for 92% of transporter expression in myeloid cells and was expressed 17-fold higher than *SLC16A1* in carcinoma. This predominance was independently reproduced in the Cillo and Kürten cohorts (27- and 22-fold, respectively), compared with only 1.4–2.4-fold in T cells across the three datasets (Fig. 5B). Among the nine major cell types, myeloid cells showed the highest *SLC16A3* and lowest *SLC16A1* expression.

**Figure 5.**
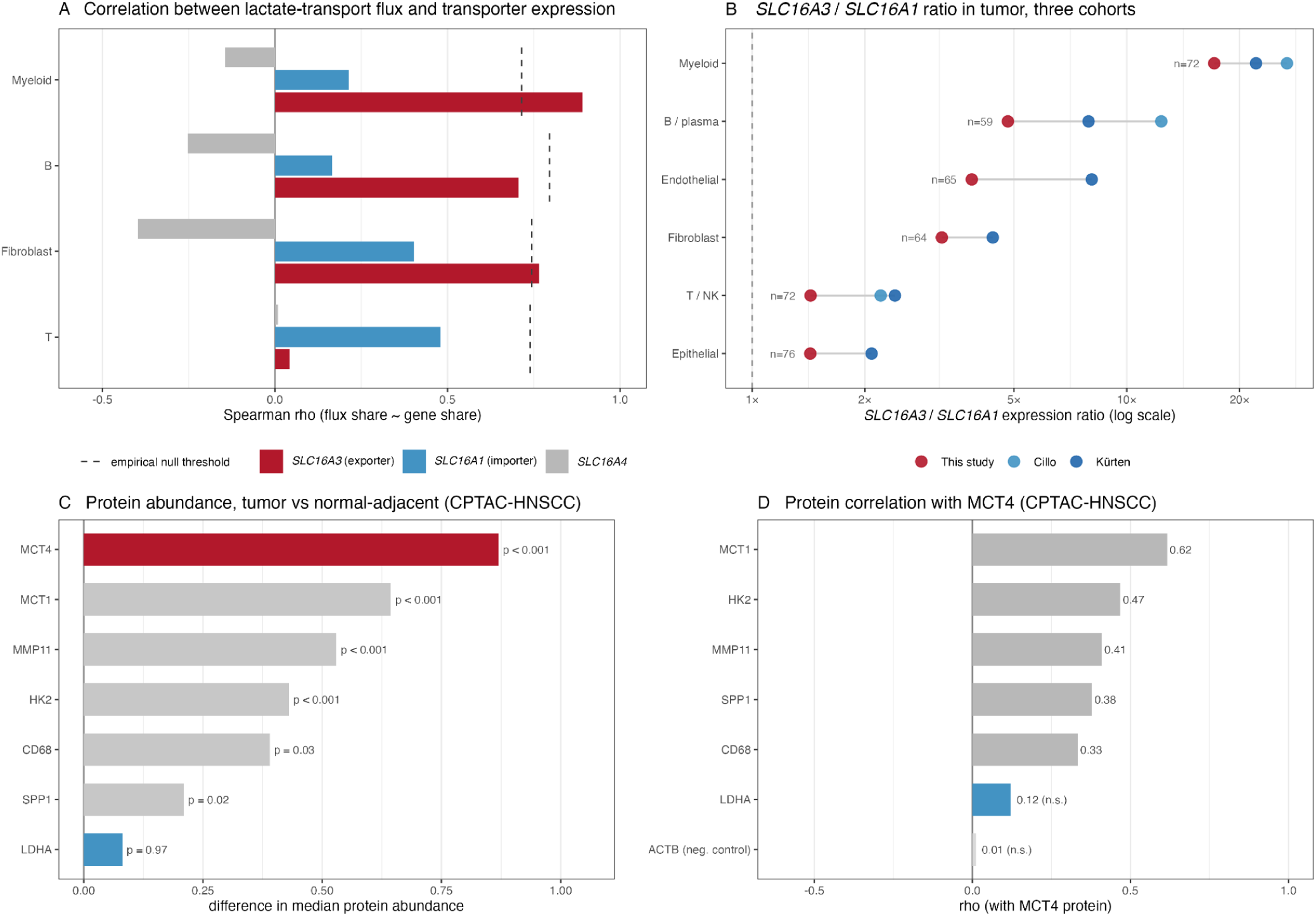
MCT4 is the dominant lactate transporter in myeloid cells. **(A)** Correlation between lactate-transport flux and transporter expression across cell types, ordered by *SLC16A3* share of total transporter expression. Dashed lines indicate the empirical null threshold for each cell type and module size. **(B)** *SLC16A3* (MCT4) to *SLC16A1* (MCT1) expression ratios across cell types in tumor tissue from three independent HNSCC datasets. Cell types with fewer than 8 samples were excluded. **(C)** Difference in median protein abundance between tumor and normal-adjacent tissue for seven proteins in CPTAC-HNSCC. LDHA (blue) is shown as a glycolytic comparator. **(D)** Correlation of MCT4 protein abundance with the indicated proteins across CPTAC-HNSCC samples. ACTB serves as a negative control.

During progression from normal mucosa to carcinoma, *SLC16A3* expression increased in all 10 paired myeloid samples (log2FC = 1.19, FDR = 8 × 10⁻⁴), whereas *SLC16A1* showed a modest, nonsignificant decrease (log2FC = −0.46, rising in 3 of 10, FDR = 0.140). This analysis was performed directly on non-imputed expression data and was independent of scFEA flux estimation.

At the protein level, CPTAC-HNSCC analysis showed strong enrichment of MCT4 in tumors relative to normal-adjacent tissues (difference in median MCT4 protein abundance = 0.87, Mann-Whitney P = 4 × 10⁻¹²; Fig. 5C). *SLC16A3* transcript and MCT4 protein abundance were highly correlated (ρ = 0.83). The MCT4 protein abundance is also correlated with MCT1, HK2, SPP1, MMP11, and the macrophage marker CD68, but not with LDHA (ρ = 0.12; Fig. 5D). Together, these data identify MCT4 as the predominant lactate transporter associated with the increased lactate-export program in myeloid cells.

### Lactate metabolic gene expression is associated with adverse clinical outcome

We next examined the clinical significance of the lactate metabolic gene program in TCGA-HNSC. Based on our above data, we constructed a three-gene program comprising *SLC16A3*, *LDHA*, and *HK2*, considering that the three genes represent key steps in lactate metabolism, encompassing glucose utilization (*HK2*), lactate production (*LDHA*), and lactate export (*SLC16A3*). Importantly, this lactate metabolic gene program was associated with worse overall survival (HR = 1.35 per standard deviation, 95% CI 1.07–1.70, P = 0.010; n = 417, 184 events; Fig. 6A). The association was stronger than that of the individual genes, including *LDHA* (HR = 1.20, P = 0.015), *SLC16A3* (HR = 1.14, P = 0.10), and *HK2* (HR = 1.08, P = 0.30; Fig. 6B).

**Figure 6.**
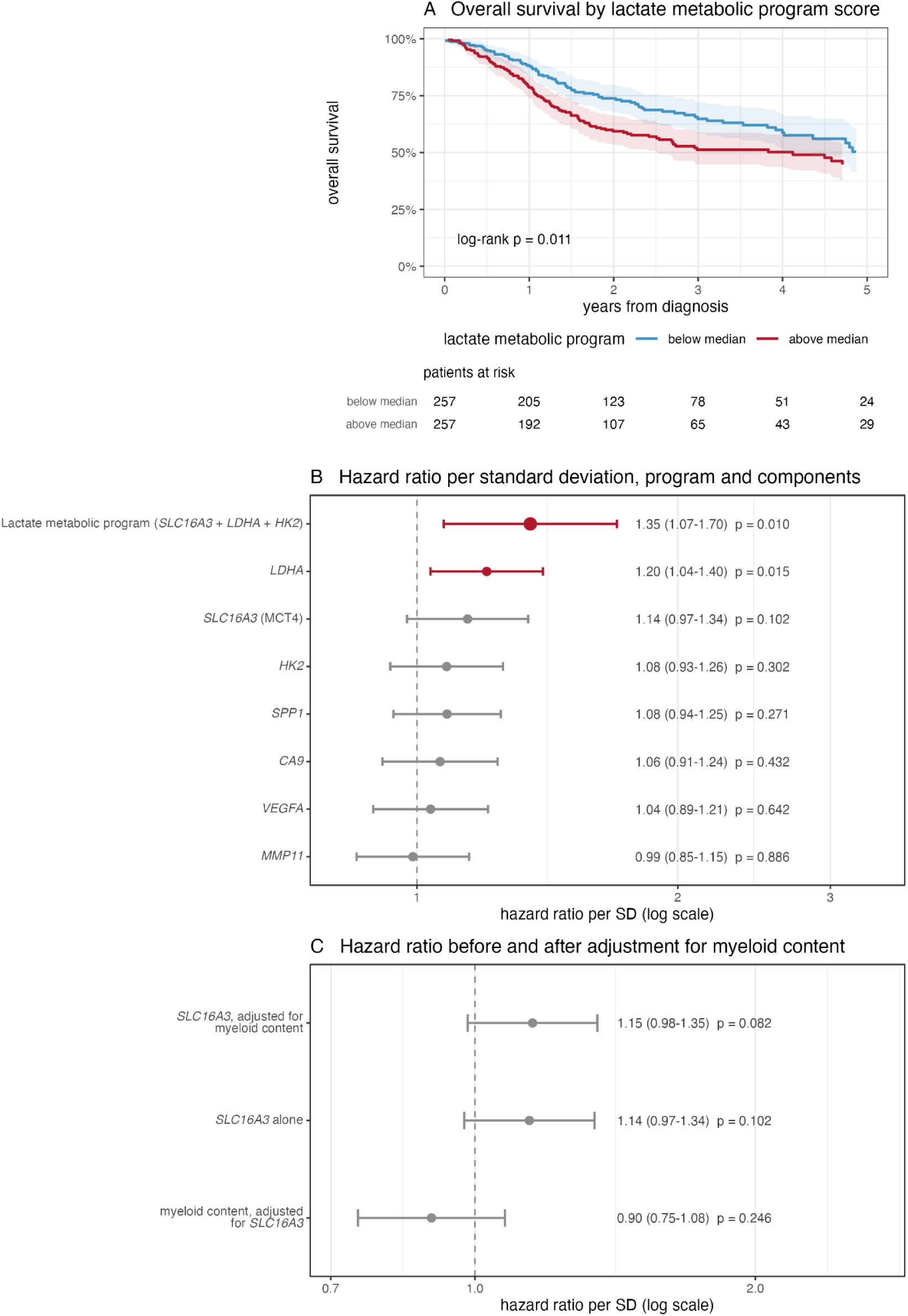
Clinical association of the lactate metabolic program in TCGA-HNSC. (**A**) Overall survival stratified by median program score (*SLC16A3*, *LDHA*, and *HK2*; log-rank P = 0.011). Numbers at risk are shown; the median split is used for visualization only. (**B**) Cox regression analysis of the program and individual genes, adjusted for age and stratified by stage (n = 417, 184 events). Hazard ratios are shown per standard deviation. (**C**) Association of *SLC16A3* with overall survival before and after adjustment for myeloid content.

Because TCGA represents bulk tumor expression, we asked whether the association with *SLC16A3* simply reflected differences in myeloid abundance. Adjustment for a myeloid content score had minimal effect on the *SLC16A3* hazard ratio (HR = 1.14 before and 1.15 after adjustment), whereas myeloid content itself was not associated with survival after accounting for *SLC16A3* (HR = 0.90, P = 0.25; Fig. 6C).

Finally, because *SLC16A3* is transcriptionally regulated by HIF-1α, we examined its relationship with tumor hypoxia. Indeed, *SLC16A3* expression correlated with three established hypoxia signatures in TCGA-HNSC (Buffa: ρ = 0.36; Winter: ρ = 0.29; Ragnum: ρ = 0.24; n = 514; Supplementary Fig. S1). In contrast, *SLC16A3* expression was not associated with HPV status, either across HNSCC (P = 0.71) or within oropharyngeal tumors (P = 0.87). Together, these findings link increased lactate metabolic gene expression to tumor hypoxia and adverse clinical outcome.

## Discussion

Our study identifies increased lactate export as a prominent feature of myeloid metabolic reprogramming during HNSCC progression. Myeloid cells showed the highest and most consistent allocation to lactate export among major cell types, and this allocation increased from normal mucosa to carcinoma in every paired patient, with the median allocation nearly doubling. This program was dominated by MCT4/*SLC16A3*, a finding reproduced across two independent HNSCC single-cell cohorts.

These findings extend previous work implicating MCT4 in oral carcinogenesis. Bisetto et al. identified MCT4 expression in CD163+ macrophages in human HNSCC and showed that MCT4 deficiency reduced invasive lesions in a 4NQO model [3]. Our data provide additional cellular resolution, showing substantial heterogeneity in lactate-export allocation across myeloid subtypes, with the highest allocation in SPP1+ macrophages. However, the increase from normal mucosa to carcinoma remained when these cells were excluded, suggesting a broader metabolic alteration across the myeloid compartment rather than an effect driven by a single tumor-associated population.

Notably, increased lactate export was not simply a consequence of globally increased glycolysis. Lactate-production allocation increased less consistently, while a TCA-cycle module increased alongside lactate export. Similarly, *SLC16A3* increased in every paired myeloid sample, whereas *SLC16A1* did not. Together, these findings suggest selective enhancement of lactate-export capacity rather than a simple shift from oxidative toward glycolytic metabolism during tumor development. The preferential association of MCT4 with myeloid cells may also extend beyond HNSCC: in glioma, *SLC16A3* has similarly been localized predominantly to immune populations, whereas *SLC16A1* is preferentially expressed by malignant cells [15].

This distinction may have therapeutic implications. Indeed, MCT4-dependent lactate secretion has been implicated in immune suppression and resistance to checkpoint blockade [16], and MCT4 inhibition can enhance immunotherapy responses in preclinical models [17]. These studies have largely focused on tumor-cell-derived lactate. Our findings raise the possibility that myeloid cells represent an additional source of MCT4-associated lactate export within the tumor microenvironment. We did not observe a robust association between myeloid lactate-export allocation and immunosuppressive transcriptional programs (data not shown); however, this does not exclude an immunoregulatory function, as exported lactate may act primarily on neighboring cells. Compartment-specific perturbation of MCT4 will be required to determine whether myeloid-derived lactate directly contributes to tumor progression or immune suppression.

We also found the lactate program has potential for carrying prognostic information. The TCGA-HNSC analysis showed an association between the three-gene program (SLC16A3, LDHA, and HK2) and survival that was not explained by differences in myeloid abundance. However, bulk expression cannot localize the signal to the myeloid compartment, and establishing this program as a biomarker will require validation in independent cohorts.

Several limitations of our study should be considered. Progression analysis was based on ten patients with paired normal mucosa and carcinoma, and independent validation in a larger paired cohort will be important. In addition, scFEA provides model-based rather than direct measurements of metabolic flux; although we applied stringent concordance and reproducibility criteria, our analysis cannot establish the absolute amount of lactate exported by each cell population. Finally, subtype-restricted analyses demonstrate that the increase is not explained solely by myeloid composition but do not establish lineage continuity between normal and tumor-associated populations. Spatially resolved MCT4 protein measurements, direct metabolic assays, and compartment-specific MCT4 perturbation will be important next steps to establish the functional significance of myeloid lactate export in HNSCC.

## Acknowledgments

This work was supported by the NIH under the award R37CA237022, R01DE033648, R01DK135562 and 1R01DE035339 (D.-C.L.), Kure It Cancer Research as well as USC Office of Research and Innovation (OORI) Research Catalyst Program.

The authors used Claude Opus 5.0 (Anthropic) to assist with proofreading, editing, and condensing of the manuscript text. Separately, Claude was used to generate, debug, and review analysis and figure code which the authors inspected and refined. The authors take full responsibility for the content of the publication.

## Supplementary Information

Supplementary Figure S1. Correlation of *SLC16A3* expression with three published hypoxia signatures in TCGA-HNSC.

Supplementary Table S1. Additional analyses not meeting predefined robustness criteria.

Supplementary Table S2. The 19 progression comparisons meeting the FDR, effect-size and cross-cohort concordance criteria, with flux–enzyme concordance status.

Supplementary Table S3. Genes differentially expressed between normal mucosa and carcinoma in myeloid, fibroblast and T cells (edgeR, FDR < 0.10).

Supplementary Table S4. HPV-stratified comparisons of lactate-handling modules (M_4, M_6, M_78) across myeloid subtypes in carcinoma, with per-arm patient counts, effect sizes, and FDR.

## References

1. Fang C, Zhou S, Yu K, Zhang L, Lian W, Tie X, et al. Lactate metabolism and lactylation in cancer: from pathogenesis to therapeutic advances. Signal Transduct Target Ther. 2026 May 21;11(1):190. doi:10.1038/s41392-026-02672-x

2. Jiang B. Aerobic glycolysis and high level of lactate in cancer metabolism and microenvironment. Genes Dis. 2017 Mar;4(1):25–7. doi:10.1016/j.gendis.2017.02.003 PubMed PMID: 30258905; PubMed Central PMCID: PMC6136593.

3. Bisetto S, Whitaker-Menezes D, Wilski NA, Tuluc M, Curry J, Zhan T, et al. Monocarboxylate Transporter 4 (MCT4) Knockout Mice Have Attenuated 4NQO Induced Carcinogenesis; A Role for MCT4 in Driving Oral Squamous Cell Cancer. Front Oncol. 2018 Aug 28;8:324. doi:10.3389/fonc.2018.00324

4. Alghamdi N, Chang W, Dang P, Lu X, Wan C, Gampala S, et al. A graph neural network model to estimate cell-wise metabolic flux using single-cell RNA-seq data. Genome Res. 2021 Oct;31(10):1867–84. doi:10.1101/gr.271205.120

5. Liu ZL, Meng XY, Bao RJ, Shen MY, Sun JJ, Chen WD, et al. Single cell deciphering of progression trajectories of the tumor ecosystem in head and neck cancer. Nat Commun. 2024 Mar 22;15(1):2595. doi:10.1038/s41467-024-46912-6

6. Choi JH, Lee BS, Jang JY, Lee YS, Kim HJ, Roh J, et al. Single-cell transcriptome profiling of the stepwise progression of head and neck cancer. Nat Commun. 2023 Feb 24;14(1):1055. doi:10.1038/s41467-023-36691-x

7. Kim S, Kee HJ, Kim D, Jang J, Jeong HO, Sim NS, et al. Multiregional single-cell transcriptomics reveals an association between partial EMT and immunosuppressive states in oral squamous cell carcinoma. iScience. 2025 Sep;28(9):112988. doi:10.1016/j.isci.2025.112988

8. Bill R, Wirapati P, Messemaker M, Roh W, Zitti B, Duval F, et al. *CXCL9:SPP1* macrophage polarity identifies a network of cellular programs that control human cancers. Science. 2023 Aug 4;381(6657):515–24. doi:10.1126/science.ade2292

9. Puram SV, Mints M, Pal A, Qi Z, Reeb A, Gelev K, et al. Cellular states are coupled to genomic and viral heterogeneity in HPV-related oropharyngeal carcinoma. Nat Genet. 2023 Apr;55(4):640–50. doi:10.1038/s41588-023-01357-3

10. Cillo AR, Kürten CHL, Tabib T, Qi Z, Onkar S, Wang T, et al. Immune Landscape of Viral- and Carcinogen-Driven Head and Neck Cancer. Immunity. 2020 Jan;52(1):183–199.e9. doi:10.1016/j.immuni.2019.11.014

11. Kürten CHL, Kulkarni A, Cillo AR, Santos PM, Roble AK, Onkar S, et al. Investigating immune and non-immune cell interactions in head and neck tumors by single-cell RNA sequencing. Nat Commun. 2021 Dec 17;12(1):7338. doi:10.1038/s41467-021-27619-4

12. Huang C, Chen L, Savage SR, Eguez RV, Dou Y, Li Y, et al. Proteogenomic insights into the biology and treatment of HPV-negative head and neck squamous cell carcinoma. Cancer Cell. 2021 Mar;39(3):361–379.e16. doi:10.1016/j.ccell.2020.12.007

13. Cerami E, Gao J, Dogrusoz U, Gross BE, Sumer SO, Aksoy BA, et al. The cBio Cancer Genomics Portal: An Open Platform for Exploring Multidimensional Cancer Genomics Data. Cancer Discov. 2012 May 1;2(5):401–4. doi:10.1158/2159-8290.CD-12-0095

14. Gao J, Aksoy BA, Dogrusoz U, Dresdner G, Gross B, Sumer SO, et al. Integrative Analysis of Complex Cancer Genomics and Clinical Profiles Using the cBioPortal. Sci Signal. 2013 Apr 2;6(269). doi:10.1126/scisignal.2004088

15. Zhu T, Ge X, Gong S, Guo S, Tao Q, Guo J, et al. Prognostic value of lactate transporter SLC16A1 and SLC16A3 as oncoimmunological biomarkers associating tumor metabolism and immune evasion in glioma. Cancer Innov. 2022 Oct;1(3):229–39. doi:10.1002/cai2.32

16. Qian Y, Galan-Cobo A, Guijarro I, Dang M, Molkentine D, Poteete A, et al. MCT4-dependent lactate secretion suppresses antitumor immunity in LKB1-deficient lung adenocarcinoma. Cancer Cell. 2023 Jul;41(7):1363–1380.e7. doi:10.1016/j.ccell.2023.05.015

17. Babl N, Decking SM, Voll F, Althammer M, Sala-Hojman A, Ferretti R, et al. MCT4 blockade increases the efficacy of immune checkpoint blockade. J Immunother Cancer. 2023 Oct;11(10):e007349. doi:10.1136/jitc-2023-007349

