## Supplementary Figure S1 for "MCT4-Associated Lactate Export Defines Myeloid Metabolic Reprogramming during Head and Neck Cancer Progression"

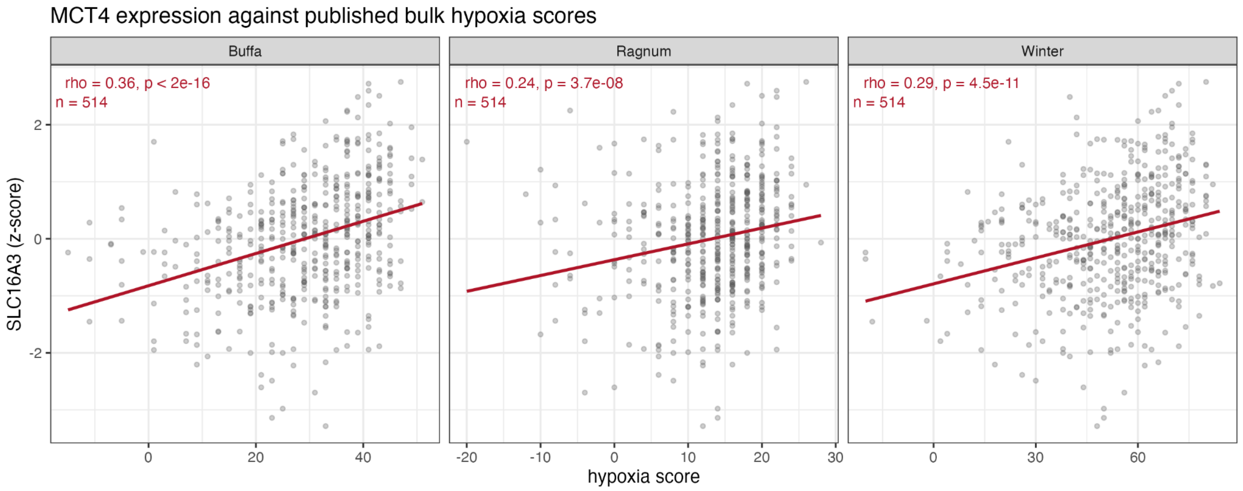


**Supplementary Figure S1. Correlation of SLC16A3 expression with three published hypoxia signatures in TCGA-HNSC.** SLC16A3 expression (z-score) against the Buffa, Ragnum, and Winter bulk hypoxia scores, one point per tumor (n = 514). Spearman ρ = 0.36 ($P < 2 \times 10^{-16}$), 0.24 ($P = 3.7 \times 10^{-8}$), and 0.29 ($P = 4.5 \times 10^{-11}$), respectively; red lines are linear fits. The single gene is shown rather than the three-gene export program because LDHA and HK2 are members of the Buffa and Winter metagenes, whereas SLC16A3 belongs to neither. These are bulk measurements and cannot attribute the association to any cell type.
